# A nonlinear simulation framework supports adjusting for age when analyzing BrainAGE

**DOI:** 10.1101/377648

**Authors:** Trang T. Le, Rayus Kuplicki, Brett A. McKinney, Hung-wen Yeh, Wesley K. Thompson, Tulsa 1000 Investigators, Martin P. Paulus

**Author notes:** **Correspondence:** Rayus Kuplicki. These authors contributed equally to this work (Trang T. Le, Rayus Kuplicki).

## Abstract

Several imaging modalities, including T1-weighted structural imaging, diffusion tensor imaging, and functional MRI can show chronological age related changes. Employing machine learning algorithms, an individual’s imaging data can predict their age with reasonable accuracy. While details vary according to modality, the general strategy is to: 1) extract image-related features, 2) build a model on a training set that uses those features to predict an individual’s age, 3) validate the model on a test dataset, producing a predicted age for each individual, 4) define the “Brain Age Gap Estimate” (BrainAGE) as the difference between an individual’s predicted age and his/her chronological age, and 5) estimate the relationship between BrainAGE and other variables of interest, and 6) make inferences about those variables and accelerated or delayed brain aging. For example, a group of individuals with overall positive BrainAGE may show signs of accelerated aging in other variables as well. There is inevitably an overestimation of the age of younger individuals and an underestimation of the age of older individuals due to ‘regression to the mean’. The correlation between chronological age and BrainAGE may significantly impact the relationship between BrainAGE and other variables of interest when they are also related to age. In this study, we examine the detectability of variable effects under different assumptions. We use empirical results from two separate datasets [training=475 healthy volunteers, aged 18 – 60 years (259 female); testing=489 participants including people with mood/anxiety, substance use, eating disorders and healthy controls, aged 18 – 56 years (312 female)] to inform simulation parameter selection. Outcomes in simulated and empirical data strongly support the proposal that models incorporating BrainAGE should include chronological age as a covariate. We propose either including age as a covariate in step 5 of the above framework, or employing a multistep procedure where age is regressed on BrainAGE prior to step 5, producing BrainAGE Residualized (BrainAGER) scores.

## 1 Introduction

Aging is a biological process that can affect behavioral and cognitive dimensions. Biological age as measured by telomere length deviates from an individual’s chronological age as a result of environment, lifestyle, and genetics (Shammas, 2011). However, other measures of biological age that may be particularly relevant to psychopathology can involve structural and functional changes in the brain.

Several imaging modalities, including T1-weighted structural imaging (Franke et al., 2010), diffusion tensor imaging (Han et al., 2014; Lin et al., 2016), and functional MRI (Tian et al., 2016) have been used in conjunction with machine learning algorithms to predict an individual’s age. Recently, integration of neuroimaging data of different feature types and across multiple modalities has been shown to improve age prediction (Erus et al., 2015; Gutierrez Becker et al., 2018; Liem et al., 2017). While the details vary according to modality, the general strategy has been to 1) extract image-related features, 2) build a model on a training set composed of healthy participants using these features to predict participant age, 3) apply that model to a testing set, producing a predicted age for each individual, 4) compute the difference between a participant’s predicted age and chronological age (often referred to as Brain Age Gap Estimate, BrainAGE, or brain predicted age difference, brain-PAD), 5) test for relationships between other variables of interest and BrainAGE, and 6) make inferences about accelerated or delayed brain aging (Cole and Franke, 2017). Variables of interest have included physical fitness (Ritchie et al., 2017), physical activity (Steffener et al., 2016), cognitive impairment after traumatic brain injury (Cole et al., 2015), mortality risk in elderly participants (Cole et al., 2018), acute ibuprofen administration in healthy participants (Le et al., 2018) or status of various diseases and disorders such as diabetes (Franke et al., 2013), Alzheimer’s disease (Gaser et al., 2013; Löwe et al., 2016), psychiatric disorders (Koutsouleris et al., 2014; Nenadić et al., 2017) and human immunodeficiency virus (Wilkins, 2017).

Support Vector Regression (SVR) with a radial kernel is a commonly used machine learning algorithm to predict age and compute BrainAGE (Franke et al., 2010), along with other methods such as Gaussian process and relevant vector regression (Drucker et al., 1997). The residual error of these age-predicting models, BrainAGE, is necessarily correlated with age, which results in an overestimation of the age of younger individuals and an underestimation of the age of older individuals. This is due to the fact that these algorithms, like all regression methods, are subject to the fundamental phenomenon of “regression towards the mean” (Galton, 1886). A theoretical basis for this phenomenon is presented in section 2.1. In practice, the correlation between chronological age and BrainAGE is visually evident in many figures of chronological versus predicted age (Cole et al., 2018; Franke et al., 2010). While most studies involving BrainAGE have not discussed the age-BrainAGE correlation, some have accounted for this correlation by using predicted age as the primary outcome, which is similar to the correction we propose (Erus et al., 2015; Habes et al., 2016).

The age-BrainAGE correlation may affect the apparent relationship between BrainAGE and variables of interest when these other variables are also related to age. In the clinical neuroscience domain, for example, we may be interested in covariates including physiological variables such as body composition or psychological measures of mood or testing performance, some of which have clear relationships with age. In this study, we examine the detectability of multiple covariate effects in both real and simulated data. Using real data, we characterized relationships between BrainAGE, age, and other variables of interest. Then, we generated a known “ground truth” with characteristics similar to what we observed in real data. In our simulation model, age has a direct effect on the variables of interest, which may in turn affect simulated imaging features. We include both linear and nonlinear effects at each level.

The goals of the current study are: 1) to highlight the universal correlation between chronological age and BrainAGE in theory and practice and 2) develop a general framework for simulating age-dependent data that can be used to investigate the effect of the age-BrainAGE correlation in subsequent analyses. One of the challenges of determining the best practices for using BrainAGE in statistical modeling is related to the fact that variables of interest may be related to age, but not directly related to accelerated or delayed brain aging. In that case, spurious relationships with BrainAGE may be observed. Our results strongly support the proposal that models including BrainAGE as an independent variable should be adjusted for chronological age as well.

## 2 Methods

We begin with a theoretical explanation for regression toward the mean and the concurrent correlation between the residuals and observed values for any regression. Then, we show in our own data the relationships between chronological age, BrainAGE, and other covariates of interest as a basis for the parameters in our simulations. Finally, we describe a simulation approach to generate data with a comparable age effect on brain image features and show how the age-BrainAGE correlation can contribute to observed relationships, even when the simulated independent variables do not associate with imaging features. The R scripts for simulation and analysis are publicly available on the GitHub repository https://github.com/lelaboratoire/BrainAGE-simulation.

### 2.1 Theoretical Basis for the age-BrainAGE Correlation

#### 2.1.1 Regression Toward the Mean

Consider *n* data points (*y*_*i*_, *x*_*i*_), *i* = 1, …, *n* used to fit a simple linear regression *y* = *α* + *βx* + *ε*. Least-square estimation leads to

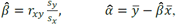

where *r*_*xy*_ is the Pearson correlation between x and y, *S*_*x*_ and *S*_*y*_ are the standard deviation, respectively. Substituting the formulas into the fitted values 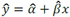 yields

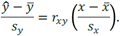

In this setting, regression toward the mean refers to the phenomenon that the standardized predicted value of y is closer to its mean than that of x to its mean for any imperfect correlation, −1 < *r*_*xy*_ < 1. The weaker the correlation, the greater the extent of regression toward the mean. For perfect correlations (|*r*_*xy*_| = 1), the standardized distance between the predicted value in *y* to its mean equals that of *x* to its mean and there is no regression toward the mean. The implication for BrainAGE is that the age of younger individuals tends to be overestimated and the age of older individuals tends to be underestimated.

#### 2.1.2 Partition of Variance or Analysis of Variance (ANOVA)

In the general setting *y* = *ƒ*(*X*) + *ϵ*, where *X* can be any dimension and *ƒ*(·) can be any regression model, the variance of *y* is partitioned into a part that can be explained by *X*, and a part due to random error: 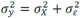. Then

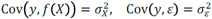

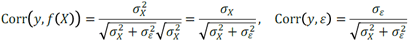

For 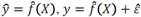 and

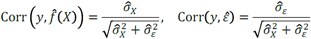

where 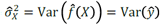 and 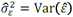.

Thus, 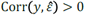 unless 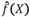 predicts y perfectly with 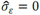. The correlation formulas suggest that the correlation between residual and *y* decreases with the correlation between *y* and 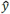, i.e. prediction accuracy of 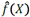. Figure S1 illustrates this phenomenon using a simple simulation where *y* was a function of x plus random normal noise. As the noise decreases (and fit increases), the correlation between *y* and the residuals decreases as well.

In the context of BrainAGE, the goal is to find 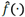 that best predicts chronological age (*y*) using brain measures as *X*, and BrainAGE is computed as 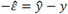. Because 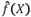 never predicts chronological age perfectly, BrainAGE remains correlated with age. When BrainAGE is used as the response variable in subsequent analyses to make inferences on a covariate *Z*, it is important to check whether *Z* is not associated with chronological age, then one may simply evaluate the bivariate association between BrainAGE and *Z*. On the other hand, if *Z* is associated with chronological age. If *Z* is associated with both chronological age and BrainAGE, chronological age may confound the relationship between BrainAGE and *Z* (Elwood, 1992) and should be taken into account. Confounding effects can be addressed at study design (e.g., randomization and matching) or in statistical analysis (e.g., stratification of the confounder or including the confounder as a covariate (Pourhoseingholi et al., 2012). For example, Franke et al. (2010) considered a variable *Z* that represents two groups (ill versus healthy) and selected two groups of individuals with similar chronological age (so *Z* is not associated with chronological age) to compare their BrainAGE. In the current work, we include chronological age as a covariate and evaluate this approach in the context of BrainAGE.

### 2.2 Empirical Data

We used two separate datasets to illustrate the correlation between BrainAGE and chronological age and the effect this can have on associations with covariates of interest. All data were collected at the Laureate Institute for Brain Research between 2009 and 2017. All protocols were approved by Western Institutional Review Board (www.wirb.com). Participants signed written informed consent and received financial compensation for their participation.

#### 2.2.1 Training Dataset

Structural MRI data were collected from 475 healthy volunteers (mean age ± sd = 30.5 ± 10.3 years; age range = 18 – 60 years; 259 female) between 2009 and 2017. Each participant was scanned in a 3T GE MR750 whole body scanner. Scans were acquired using axial T1-weighted MP-RAGE sequences with a 24cm FOV, 256×256 acquisition matrix, 8-degree flip angle and .9375x.9375mm in-plane resolution with no gap. Other parameters varied within the following ranges: 5.736 to 6.292ms TR, 1.896 to 2.104ms TE, 0.9 to 1.2mm slice thickness, with either an 8- (General Electric, Milwaukee, WI) or 32- (Nova Medical Inc., Wilmington MA) channel phased array coil. Healthy neuropsychiatric status was assessed using either the MINI-international Neuropsychiatric Interview (Sheehan et al., 1998) or the Structured Clinical Interview for DSM-IV (First et al., 2002) (First, Michael B., Spitzer, Robert L, Gibbon Miriam, and Williams, Janet B.W., 2002).

#### 2.2.2 Testing Dataset

Structural MRI data were collected from 489 (mean age ± sd = 34.6 ± 10.6 years; age range = 18 – 56 years; 312 female) participants as part of Tulsa 1000, a longitudinal observational study including people with mood/anxiety, substance use, eating disorders and healthy controls. Inclusion criteria for the participant populations were Patient Health Questionnaire ≥ 10, Overall Anxiety Severity and Impairment Scale ≥8, Drug Abuse Screening Test > 3, or SCOFF ≥ 2. Exclusion criteria included a history of significant brain trauma, neurological disorders, change in medication within six week prior to scanning, bipolar disorder, and schizophrenia. Scanning parameters for this dataset were: 24cm FOV, 256×256 acquisition matrix, 186 axial slices, 0.9mm slice thickness with no gap, TR/TE=5/2.012ms, using an 8-channel phased array coil (General Electric, Milwaukee, WI). Testing and training sets differed on mean age (t = 6.2, p < 0.0001, mean difference 4.2 years) and sex composition (χ^2^ = 8.2, p = 0.004).

All participants in the testing dataset also underwent an intense battery of assessments including self-report, clinical interviews, neuropsychological testing, and body composition analysis. For full details, please see (Victor et al., 2018). From these, we selected 154 measures, which were used to illustrate the normal range of correlations with age and how these can affect the relationship between BrainAGE and covariates of interest.

#### 2.2.3 Image Processing

All images in both the testing and training sets were processed using Freesurfer version 6.0.0 (Dale et al., 1999) in order to produce grey/non-grey matter masks. Then, using a procedure similar to Franke (Franke et al., 2010) but implemented in AFNI, all grey matter masks were transformed to MNI space via affine transformation, smoothed with an 8mm gaussian kernel, and downsampled to 8×8×8mm voxels. This produced a set of 3707 voxels per participant, with the value at each voxel representing the fraction of that voxel comprised of grey matter.

R (version 3.2.2) and R package caret (version 6.0.76) were used to fit a support vector regression (SVR) model with radial basis functions. The *ε* (tolerance margin) was fixed at and cost parameters were tuned using 5 repeats of 10-fold cross validation in the training set. The hyperparameter space was sampled using a grid search that fixed *ε* at 0.000145 and allowed cost to vary from 0.25 to 4096. The final best model (cost = 2) was then applied to the testing set to produce one predicted age for each participant. BrainAGE was taken to be predicted age minus chronological age.

Additionally, we define the Brain Age Gap Estimate Residualized (BrainAGER) to be the residual of the regression of BrainAGE on age to remove the remaining linear bias of age. This way, we have a measure of deviation from expected age that is linearly uncorrelated with chronological age.

### 2.3 Simulation

To investigate the effect of the age-BrainAGE correlation on subsequent modeling results, we simulated hierarchical correlation structures among brain features, chronological age and covariates using a generative biological model (Fig. 1). We then generated two groups of independent variables. Within each group of variables, some are dependent on age and others are not. One group was used in the simulation of neuroimaging features, while the other was not. We randomly split the data set into two subsets, trained SVR on the training set and computed BrainAGE on the testing set. On the testing set, we conducted linear regressions of BrainAGE on all independent variables, both with and without chronological age. With 1,000 replications, we assessed the significance of the contribution from the independent variables by examining the distribution of the resulting *p*-values.

**Figure 1.**
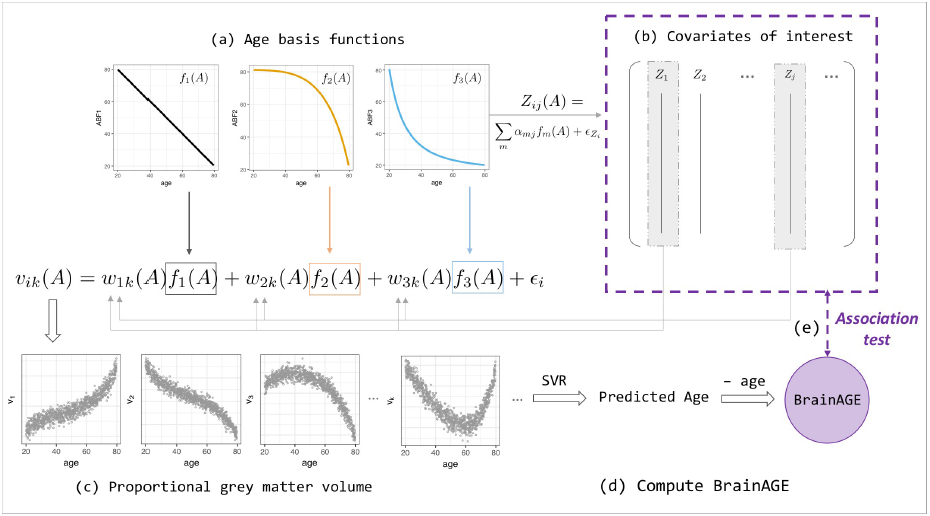
BrainAGE simulation and analysis framework (Eqs. 2-4). (a) Linear and non-linear age basis functions (ABFs) *ƒ*_*i*_ (**orange**, **black** and **blue** lines). For a particular individual *i*, the ABFs are combined to create volume *k*’s grey matter proportion *ν*_*ik*_ (**orange**, **black** and **blue** arrows) and age-dependent covariates of interest, *Z*_*ij*_(*A*) with a different set of coefficients *α*_*j*_. (b) Some of the *Z*_*ij*_ are then fed back into the *w*_*ik*_ when generating volume *ν*_*ik*_, which leads to two levels of age association between covariate and BrainAGE. (c) Proportional grey matter volume (volumetric data) generated from non-linear combinations of ABFs. (d) Predicted-age and BrainAGE computed from simulated volumetric data and simulated chronological age with Support Vector-based regression; (e) Test for association between BrainAGE and covariates of interest.

#### 2.3.1 Model Definition

A realistic simulation model should capture the properties of normal age-related brain volumetric data, such as brain region-dependent changes and nonlinear chronological age dependence (Fjell et al., 2013). A realistic simulation should also include the ability to generate age-dependent deviations from the normal population and age-dependent covariates that may influence BrainAGE nonlinearly. We consider a biological causal path model and develop a novel age-basis-function approach for simulating BrainAGE data with covariates (Fig. 1, Fig. S2).

Denoting age by *A*, we assumed an underlying (unobserved) biological process represented by *m*, which we referred to as age basis functions (ABFs). Here, without a function space defined, the term “basis” is used loosely to indicate the elementary functions that can be combined linearly to form any variable of interest y:

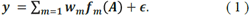

In this study, we implemented three monotone decreasing ABFs that can generate a wide range of non-linear functions (Fig. S3), and used these ABFs to simulate covariates of interest and the features extracted from an imaging modality.

##### Simulating covariates

A covariate of interest *Z*_*j*_ for participant *i* with chronological age *A*_*i*_ was generated by

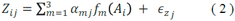

where *α*_*mj*_ is a covariate-specific weight and the covariate-specific error 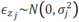 denotes a Gaussian noise with mean 0 and standard deviation *σ*_*j*_.

##### Simulating imaging modality

The proportional grey-matter volume for voxel *k* of a participant *i* with chronological age *A*_*i*_ was generated by

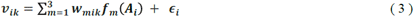

or, in short, *ν*_*ik*_ = *ƒ*(*A*_*i*_) + *∈*_*i*_, where *∈*_*i*_ represents Gaussian noise with mean 0 and standard deviation *σ*_*ν*_. This setting allows capturing within-participant correlations (4b) and spatial dependence within participants (4c):

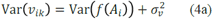

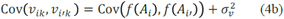

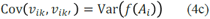

Note that the weight function *w*_*mik*_(*A*_*i*_) allows the weights of ABFs to vary across individuals and volumes, and as a function of an individual’s chronological age.

To further make the imaging modality dependent on some covariates, we let

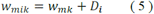

where *w*_*mk*_ is the population mean weight for ABF *ƒ*_*m*_ at voxel *k*, and the participant level departures ***D***_*i*_ depends on the first *q* variables (covariates):

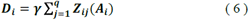

Other measurable variables, Z_j>q_ do not contribute to the weights deviation. In addition to the age-related imaging features that are generated from the ABFs, we also added 25% “background” features that do not correlate with age. Other parameters such as standard deviation of the noise *∈* were chosen with the objective of yielding *R^2^* and MAE values that closely match our empirical results when the volumetric features were used as inputs to the support vector regression (SVR) model to estimate chronological age. Nevertheless, the choice of parameters and even the simulation design matrix do not affect the overall improvement in the regression that includes age as an explanatory variable from the regression without age.

Finally, we carried out linear regressions of the covariates of interest on BrainAGE, with and without including age as an explanatory variable in the model. Over 100 replications, we assessed the detectability of the covariates as significant contributors to BrainAGE by examining their p-value distributions. In the ideal case, we should detect relationships between BrainAGE and covariates *Z*_*j*_′S.

#### 2.3.2 Simulation steps

1. Draw 1,000 age values from the uniform distribution *U*(20, 80)
2. For each *m* = 1, 2, 3, draw 100 *w*_*mik*_ values from N(0, *σ*_*w*_) for each region *k*.
3. Set *α*_*mj*_ = 0 for some m and j (Table S1). Randomly draw the remaining *α*_*mj*_ from the uniform distribution *U*(–2, –1) to construct the *j* covariate for each participant *i* (Eq. 4).
4. Construct the volumetric data set. For each imaging feature *k* of participant *i* (Eq. 2), add noisy volumetric features that do not correlate with age.
5. Randomly apply 50% of the (age, volumetric) data for training and 50% for validation. Train the SVR model using the R package `e1071` with hyperparameters set as default on the training set and apply the model on the validation set to compute the BrainAGE scores.
6. On the testing set, run linear regressions of BrainAGE on all covariates, with and without age.
7. Assess the significance of the covariates by looking at the confidence intervals of their coefficients as well as the distribution of the resulting p-values.

In steps 3 and 4, we simulated 16 covariate types in each of 1000 replicate data sets (Table S1). The 16 variables were simulated by using all 8 possible combinations of the three age basis functions. Half of them contributed to the weights *w*_*mik*_ (*A*), which consequently affected the grey matter density. For example, *Z*_*2*_ and *Z*_*10*_ were both derived from only the linear basis function *ƒ*_*1*_, but Z_10_ does not influence the aging.

Additionally, the complete simulation procedure was carried out for two scenarios: one with relatively large and another with relatively small effects of the covariates on BrainAGE. This was achieved by modifying the constant *γ* in Eq. (3) so that, in one case, the final weights *w*_*mik*_ have a larger fold change on the original weights. In particular, the fold change is computed as

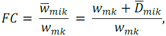

where 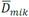 is the average of *D*_*mik*_(*A*) across all ages.

## 3 Results

### 3.1 Empirical

#### 3.1.1 Covariate Correlations with Age

Observed Pearson correlations between age and the 154 clinical variables ranged from -0.33 (PROMIS physical function) to 0.29 (waist circumference) (Fig. S4). Because any confounding effect of the correlation between age and covariates of interest is likely to be worse with larger correlations, we focused on simulated covariates that correlated with age with an r of up to 0.3.

#### 3.1.2 Age Prediction Accuracy and Bias

After fitting on the training dataset, SVR achieved a mean absolute error of 4.84 years and explained 64% of the variance in age in the testing dataset (Fig. 2a). This is comparable to the cross-validated performance on the training set, where MAE was 5.1 years and R^2^ was 0.59. The correlation between age and predicted age was 0.82. On the other hand, regression towards the mean lead to a negative relationship between age and BrainAGE (r = -0.63, Fig. 2c). After removing the linear trend as shown in Figure 2c, we observed no relationship between age and BrainAGER (r = 0.001, Fig. 2e). More explicitly, BrainAGE had a positive expected value at low chronological age and a negative expected value at high chronological age, while BrainAGER has an expected value of 0 regardless of actual age.

**Figure 2.**
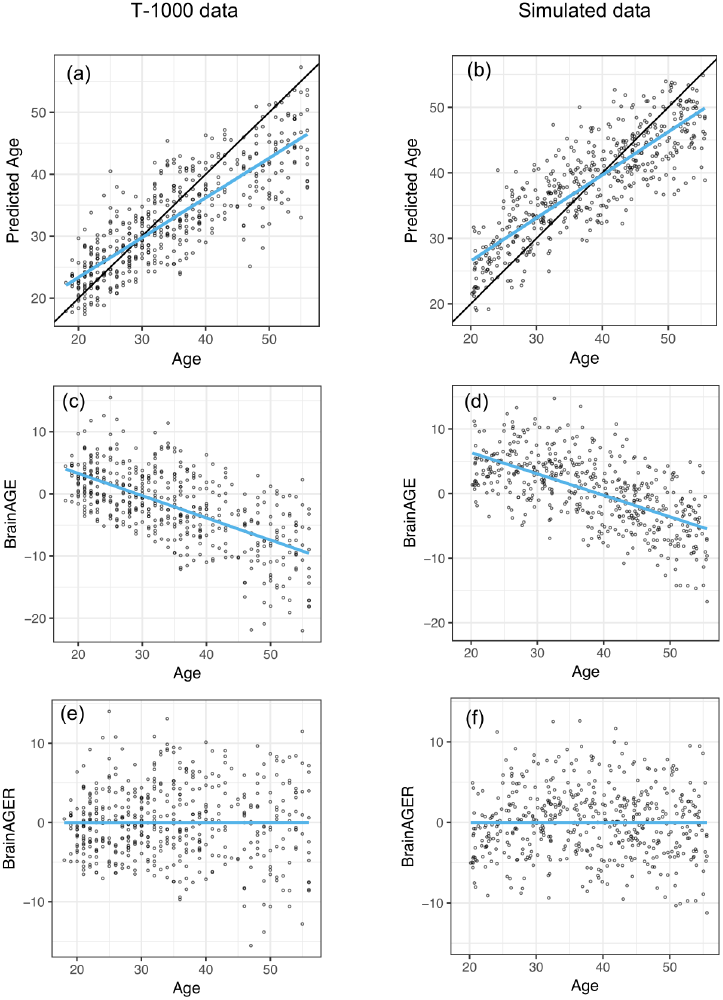
Similar out-of-sample R^2^ when applying SVR to predict age as well as negative correlation between BrainAGE and chronological age between T1000 data and simulated data. (a-b) Chronological age versus predicted age in the testing dataset, with a mean absolute error (MAE) of 4.78 years and R^2^ = 0.65 in (a) and MAE = 5.15, R^2^ = 0.841 in (b). Overlaying **black** 45-degree line and **blue** regression line showed regression toward the mean. (c-d) Chronological age versus BrainAGE (r=-0.63). Negative correlation between BrainAGE and chronological age indicates younger participants tend to have positive BrainAGE and old participants tend to have negative BrainAGE. (e-f) After removing the linear trend in b-c, there is no relationship between age and BrainAGER (r = 0.001). BrainAGER has an expected value of 0, regardless of chronological age.

#### 3.1.3 Relationships among age-covariate, covariate-BrainAGE, and covariate-BrainAGER correlations

In order to investigate the effect that the correlation between BrainAGE and chronological age can have on the conclusions of an imaging study, we computed the correlations between each of the covariates and age, BrainAGE and BrainAGER. Larger age-covariate correlations lead to larger differences in measured correlation between that covariate and BrainAGER or BrainAGE (Fig. 3a, colored points far from the 45° line). When age did not correlate with a covariate, BrainAGE and BrainAGER tended to give similar results (grey points, near the 45° line). When age positively correlated with covariates (*e.g.*, BMI), BrainAGER gave more positive values, and when age negatively correlated with covariates (*e.g.*, PROMIS physical function), BrainAGER yields more negative values. Similarly, the greater the variance explained by age, the greater the squared difference in r between using BrainAGE or BrainAGER (3b).

**Figure 3.**
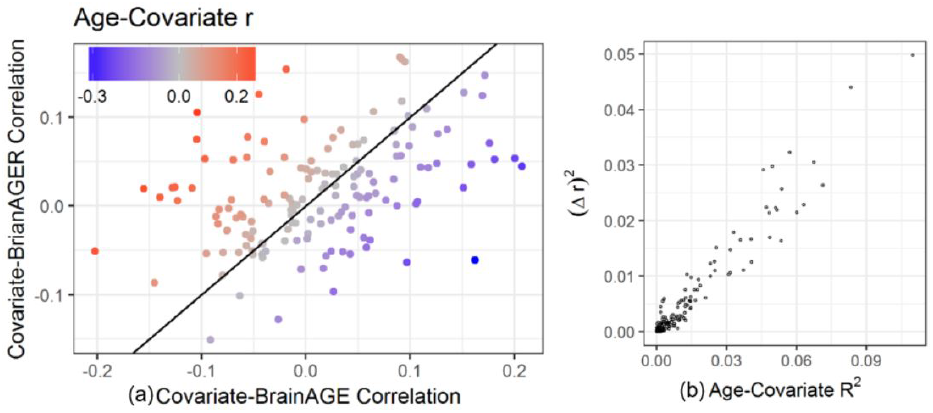
Relationship between age-covariate correlation and the difference in measured correlation. The difference between using BrainAGE and BrainAGER depends on the age-covariate relationship. (a) Covariate-BrainAGER correlations as a function of the covariate-BrainAGE correlation, with points colored according to the Age-Covariate correlation. The 45-degree line is shown, and covariates more strongly related to age are further from the line. (b) The squared difference in r between using BrainAGE and BrainAGER as a function of the variance explained by age.

Table 1 shows the top 22 variables that are significantly correlated with either BrainAGE or BrainAGER after FDR correction for 154 tests. Notably, 18 variables were related to BrainAGE, and the strongest relationships were among variables strongly correlated with age, including body composition (percent body fat r = -0.2, percent body water r = 0.2, percent dry lean mass r = 0.2) and sensation seeking (r = 0.18). BrainAGER was only significantly correlated with six variables including waist to hip ratio (r = 0.15), color naming scaled (r = -0.15), and lean body mass (r = 0.17).

**Table 1.** Correlation and significance after FDR adjustment of each covariate with BrainAGE (*r*_*BrainAGE*_, *p*_*BrainAGE*_) or BrainAGER (*r*_*BrainAGER*_, *p*_*BrainAGER*_). The last column contains the direct correlation between each covariate and age (*r*_*age*_). For brainAGE, where age is not adjusted, there are 17 covariates with FDR adjusted p-values <.05 and for BrainAGER, which residualizes age, there are six covariates with adjusted p-value < .05. Cells with p less than 0.05 are bold.

| | $r_{BrainAGE}$ | $p_{BrainAGE}$ | $r_{BrainAGER}$ | $p_{BrainAGER}$ | $r_{age}$ |
| --- | --- | --- | --- | --- | --- |
| *PROMIS_PainInterfTscore* | -0.128 | **0.047** | 0.021 | 0.91 | 0.227 |
| *PhysFunc* | 0.162 | **0.006** | -0.061 | 0.655 | -0.331 |
| *BAS_FunSeeking* | 0.159 | **0.007** | 0.047 | 0.7 | -0.201 |
| *TES_TotalOccurrence* | -0.14 | **0.025** | 0.01 | 0.971 | 0.226 |
| *IRI_EmpaConcern* | -0.145 | **0.019** | -0.086 | 0.416 | 0.11 |
| *IntSexAct* | 0.151 | **0.011** | 0.021 | 0.91 | -0.22 |
| *UPPSP_SensSeek* | 0.181 | **0.002** | 0.053 | 0.655 | -0.231 |
| *CDDR_PosReinforcement* | 0.151 | **0.047** | 0.128 | 0.238 | -0.073 |
| *PROMIS_AlcoholNegConsqTscore* | 0.172 | **0.004** | 0.147 | **0.03** | -0.095 |
| *PROMIS_AlcoholPosConsqTscore* | 0.176 | **0.003** | 0.071 | 0.545 | -0.193 |
| *PROMIS_AlcoholPosExpectTscore* | 0.127 | **0.047** | 0.081 | 0.455 | -0.098 |
| *PROMIS_AlcoUseTscore* | 0.169 | **0.004** | 0.124 | 0.108 | -0.112 |
| *DryLeanMass* | 0.095 | 0.17 | 0.162 | **0.017** | 0.042 |
| *FatMass* | -0.155 | **0.009** | 0.02 | 0.91 | 0.26 |
| *LeanBodyMass* | 0.091 | 0.183 | 0.166 | **0.017** | 0.052 |
| *PercentBodyFat* | -0.202 | **<0.001** | -0.051 | 0.655 | 0.251 |
| *Water* | 0.09 | 0.191 | 0.167 | **0.017** | 0.056 |
| *W.HRatio* | -0.019 | 0.834 | 0.154 | **0.03** | 0.223 |
| *PercentWater* | 0.2 | **<0.001** | 0.054 | 0.655 | -0.245 |
| *PercentDryLean* | 0.207 | **<0.001** | 0.044 | 0.727 | -0.267 |
| *CW_ColorNamingScaled* | -0.092 | 0.196 | -0.151 | **0.03** | -0.041 |
| *CW_InhibitionVsColorNamingScaled* | 0.135 | **0.04** | 0.086 | 0.416 | -0.108 |

### 3.2 Simulation

#### 3.2.1 Negative correlation between BrainAGE and chronological age in simulated MRI data

We set the parameters of our simulation algorithm to achieve realistic characteristics of experimental data, such as correlation distribution between volumes and chronological age and the negative correlation between computed BrainAGE and chronological age. This negative correlation was also present in previous models such as with Gaussian Process Regression (Cole et al., 2017) and Relevant Vector Regression (Franke et al., 2010). Simulated results closely mirrored empirical results. The simulated testing data had MAE of 4.58 years and R^2^ of 0.71 (2b). In our simulated data, we observed an overestimation of younger participant’s ages and an underestimation of older participant’s ages (Fig. 2d). After removing the effect of age on BrainAGE, simulated BrainAGER had an expected value of 0 regardless of actual age (2f).

#### 3.2.2 Reduction of false discoveries in regression that include age as explanatory variable

In the linear models regressing BrainAGE on the 16 covariates of interest with simulated large effect sizes (FC = 1.255), we observed the following: when age was not included as an explanatory variable, many age-related covariates were shown to have statistically significant association with BrainAGE (Fig. 4a, c), even when they did not contribute to the weights that made up the neuroimaging features (Fig. 4, **orange** boxplots above the horizontal). These false positives (FP) were simply the result of the relationship between these covariates and chronological age that are part of the BrainAGE’s defining formula. Moreover, several covariates that were simulated to contribute to the brain structure volumes had p-values on average above 0.05 (Fig 4, **blue** boxplots below the horizontal).

**Figure 4.**
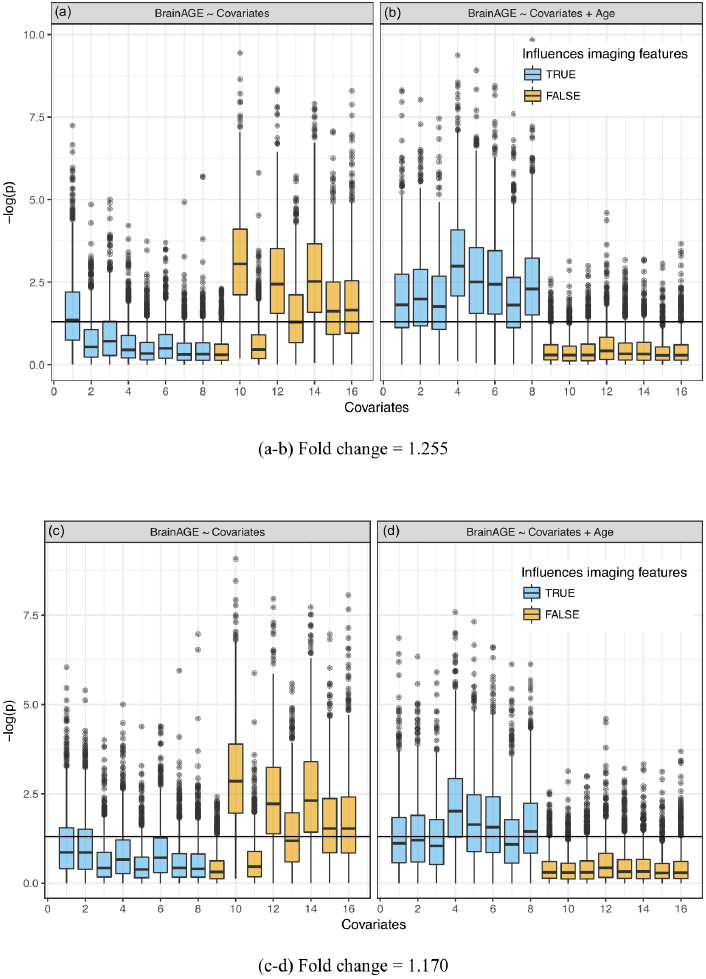
Significance of linear regression of covariates with BrainAGE for 100 replicate simulations. Each data set contains 16 age-dependent covariates with differing age dependencies (linear and nonlinear) and effects on volumetric variation. **Blue** boxes are variables that have a direct (TRUE) effect on BrainAGE, **orange** boxes are variables that do not have a direct effect on BrainAGE (FALSE), and this effect is relatively large in the top (a, b) and small in the bottom (c, d) plots. Boxplots on the left (a, c)do not use age as an explanatory variable and models on the right (b, d) include age as an explanatory variable. “Significance” was measured by –log(p). Horizontal line is at –log(0.05).

When age was included in the regression as an extra explanatory variable, the significance increased (p-values decreased) for all variables that were generated to have an association with the imaging features, even variables that were already detected in the previous regression without age (Fig. 4b, d). Further, the decrease in significance (increase in p-values) for unrelated covariates indicated a significant decrease in the number of false positives. Variation in the p-values across covariates came from their different (linear and nonlinear) age dependencies and effects on volumetric variation. In other words, the real “significance” of a covariate depended on from which age basis functions it was generated and how it affected the brain features (*w*_*1k*_, *w*_*2k*_ or *w*_*3k*_). Simulations with a smaller effect size (FC = 1.170, Fig 4c, d) showed a similar effect, though attenuated, for covariates that were contributors to *w*_*mk*_. The positive rate (true and false) across 100 replications is quantified in Supplementary Table S2. Values in this table represent the portion of each boxplot above the horizontal line, which is the TP rate for covariates that had an influence on imaging features and FP rates for covariates that did not.

## 4 Discussion

This study aims to highlight the relationship between chronological age and BrainAGE and its transitive effect on the relationship between BrainAGE and covariates of interest that are also related to age. We propose a solution to this problem: either use BrainAGER, or in the simple case of post-hoc linear regression, use chronological age as a covariate in subsequent analyses. We developed a simulation framework to generate data with complex, but known, relationships between the original imaging features, age, and a set of covariates that may also be related to age. Then, we were able to quantify the effect that accounting for age has on the ability to detect actual and spurious correlations with covariates in subsequent analyses.

Our main findings can be separated into three parts: analytical, empirical, and simulated data results. The analytical results provide a theoretical basis for the age-BrainAGE correlation, and the analyses using real and simulated data demonstrate this effect in practice. For the empirical data, there were three main findings: 1) many variables that may be of interest are correlated with age with Pearson coefficients of up to r = 0.3, 2) BrainAGE is strongly negatively correlated with chronological age (r = -0.63 in our dataset), 3) BrainAGER provides a measure of deviation between predicted and actual age that is not dependent on age, and has substantially different correlations with covariates that are correlated with age when compared to BrainAGE.

Since it is unknown which covariates are actually related to premature aging, we then developed a simulation framework to generate synthetic data. Simulated data showed: 1) similar characteristics to actual data when used to train and test a model on separate datasets, and 2) increased detectability of true positives and decreased occurrence of false positives when accounting for the age-covariate relationship, with this being modulated by the size of the simulated effect on physiology.

Based on our observations in both real and simulated data, we recommend that the relationship between chronological age and BrainAGE should be accounted for. The two methods proposed in this study are either: 1) regress age on BrainAGE, producing BrainAGER, which is centered on 0 regardless of a participant’s actual age or 2) include age as a regressor when doing follow-up analyses. In fact, these two methods will produce the same coefficients in the case of linear regression, with slightly larger t-statistics in the second case. The advantage of using BrainAGER is simplicity and generalizability; it could be used as the dependent variable in any arbitrary model, rather than being confined to simple linear regression. While the focus of this study is not to show specific correlates of premature aging, it is worth noting that 17 variables significantly correlated to BrainAGE whereas only 6 were related to BrainAGER, with 1 variable (PROMIS Alcohol Negative Consequences) overlapping between the two sets (Table 1). Thus, accounting for the age-BrainAGE relationship results in a vastly different set of positive findings and would lead to a remarkably different interpretation of these data. More explicitly, not correcting the age-BrainAGE correlation would lead to an extensive set of spurious results in this dataset.

### Limitations

There are a few cases where the age-BrainAGE correlation is not relevant. When comparing two groups with matched age, any differences in BrainAGE are not likely to be caused by the relationship with age. When the individuals being examined are in a restricted age range, there is not likely to be much contribution from the age-BrainAGE correlation. Also, when the variable of interest is not related to age, removing the effect of age makes almost no difference (Fig 3b). However, when these cases are not true, our findings suggest that we should include age as an explanatory variable in a final model that aims to detect association of brain anomalies with covariates of interest.

The magnitude of the age-BrainAGE correlation is directly related to the accuracy of the prediction model. The fact that the residuals are correlated with observed values is a characteristic of regression in general, regardless of the specific data domain, and has a theoretical basis described in section 2.1. Several factors may decrease the model performance on our testing set, and thereby increase the age-BrainAGE correlation. Specifically, the distribution of age ranges in our samples is non-uniform, which may lead to more weight being given to the middle of the distribution. There are substantial differences between the testing and training sets we used including age, sex, and diagnosis. It may therefore be possible to improve model performance on the testing set by subsampling the training set to have a more uniform distribution of ages and to match the testing set on several factors. However, model performance is already comparable across testing and training sets (R^2^ of 0.59 and MAE of 5.1 years, compared to 0.64 and 4.84) and is comparable with what has been previously reported.

Although the simulation was carefully designed and executed, because of the model’s complexity, we have not fully explored all scenarios with different simulation parameters. However, we have identified effect size as the most important parameter and showed how it influenced the results. When varying other parameters, we still observed a reduction in the number of false positives when age is included as an explanatory variable in the final regression (results not shown). Moreover, while determining the parameters, we aimed to obtain realistic patterns as we observed in real data, such as similar distributions of the correlation values.

By constructing and studying an appropriate generative model containing covariates that have linear and non-linear relationship with age, we demonstrated that the correlation between covariates and age should be considered when making inferences about the relationship between BrainAGE and these covariates.

## 6 Author Contributions

TL, RK, BM, HY, WT and MP contributed to the design of the study. TL and RK wrote the manuscript. TL performed simulation analyses and RK performed empirical analyses. All authors revised the manuscript critically for important intellectual content and approved the final manuscript.

## 8 Acknowledgments

